# Rates of female mouse ultrasonic vocalizations are low and are not modulated by estrous state during interactions with muted males

**DOI:** 10.1101/2024.11.25.625177

**Authors:** Cassidy A. Malone, Patryk Ziobro, Julia Khinno, Katherine A. Tschida

## Abstract

Vocalizations produced during courtship can communicate signaler attributes and facilitate mating success. Adult male mice produce high rates of ultrasonic vocalizations (USVs) during courtship interactions with females. For many decades, it was thought that only males produced courtship USVs, but recent studies using microphone arrays to assign USVs to individual signalers report that female mice produce a portion (5-18%) of total courtship USVs. The factors that regulate courtship USV production by female mice are poorly understood. In this study, we tested the idea that female courtship USV production is regulated by estrous state. To facilitate the detection of female USVs, we paired female subjects with males that were muted for USV production via ablation of midbrain neurons that are required for USV production but not for male sexual motivation. We report that total USVs recorded during interactions between group-housed B6 background female mice and muted males are low and are not modulated by female estrous state. Similar results were obtained for single-housed B6 background females and for single-housed outbred wild-derived female mice paired with muted males. These findings suggest either that female mice produce substantial rates of courtship USVs only when interacting with vocal male partners or that prior studies have overestimated female courtship USV production. Studies employing methods that can unambiguously assign USVs to individual signalers, regardless of inter-mouse distances, are needed to distinguish between these possibilities.

## Introduction

Vocalizations are important for communication, and many species produce vocalizations during courtship. Courtship vocalizations can communicate a variety of signaler attributes, including sex, species identity, individual identity, sexual motivation/receptivity, and fitness^1–4^. Courtship vocalizations can in turn affect the behavior and physiology of potential mates. For example, courtship vocalizations can promote proximity between mating partners^5^ and can promote reproductive physiology and courtship behaviors in potential mates^6–8^. Although both males and females in many species (including examples in rodents, primates, birds, frogs, and insects) produce courtship vocalizations, female courtship vocalizations have been comparatively less studied than those produced by males.

Adult male mice (*Mus musculus*) produce high rates of ultrasonic vocalizations (USVs) during interactions with female mice^9–14^, and these courtship USVs are produced in coordination with non-vocal courtship behaviors, including sniffing, following, and mounting^10,13–15^. For many decades, it was thought that only male mice produced USVs during courtship interactions.

Support for this idea came from observations that high rates of USVs are detected when male mice interact with anesthetized females^16^ or with females that have been devocalized via transection of the recurrent laryngeal nerve^14,17^. Male mice also produce high rates of USVs during olfactory investigation of female urine^18,19^. In contrast, only low rates of USVs are observed when female mice interact with anesthetized males^16,20^, devocalized males^14,17^, or male urine^20,21^. It is important to note that female mice are capable of producing USVs and in fact produce high rates of USVs during same-sex interactions^20,22–27^. Thus, one possibility is that female mice produce USVs during courtship interactions but do so at lower rates relative to males. This idea has been difficult to test, however, because male and female mice produce USVs that are similar in acoustic features^28,29^ and because mice do not display postural changes associated with USV production, complicating the assignment of individual USV syllables to a given mouse within a social group.

To tackle these challenges, a series of recent studies have developed and optimized microphone array-based and/or acoustic camera-based recording systems and computational approaches to localize USVs and to assign USV syllables to a single mouse within a socially interacting pair or group of mice^30–35^. These studies have estimated that females produce between 5-18% of the total USVs recorded during opposite-sex interactions. The earliest of these pivotal studies reported that female mice produce USVs when being chased by males^32^, although more recent work concluded that the majority of female USVs occur during snout-to-snout interactions with males, a scenario in which accurate localization is most challenging^35^. Together, these studies suggest that although male mice produce the majority of courtship USVs, female mice also produce USVs during courtship interactions. This idea is also bolstered by work in wild-derived mice showing that females interacting with males across a perforated barrier produce USVs^27,36^. However, the factors that regulate the production of courtship USVs by female mice remain poorly understood.

One attractive idea is that the production of female courtship USVs is regulated by sexual receptivity. Indeed, in some rodent species, females produce courtship USVs that are regulated by estrous state. For example, female hamsters produce USVs during and following courtship interactions with males, as well as during olfactory investigation of male odors and in response to playback of male USVs^7,37,38^. Female hamsters produce higher rates of courtship USVs when they are sexually receptive compared to times when they are non-receptive ^37^, and rates of female hamster USVs are regulated by sex hormones^39^. Similarly, female rats produce higher rates of courtship USVs when they are sexually receptive than when they are non-receptive^40^. However, the idea that sexual receptivity regulates courtship USV production by female mice remains untested. Some past studies included only female mice that were sexually receptive (i.e., in proestrus or estrus)^14,21,32,34,41,42^, and other studies did not track female estrous state^16,17,20,27,30,31,33,35,36^. Thus, the question of whether estrous state regulates female mouse courtship USV production remains unresolved.

In the current study, we tested the hypothesis that sexual receptivity promotes the production of courtship USVs by female mice. To facilitate the detection of USVs produced by females, we paired female subjects with male mice that were muted for USV production using a viral-genetic approach to ablate midbrain periaqueductal gray (PAG) neurons that are required for USV production, which selectively and dramatically reduces male USV production without affecting rates of male non-vocal courtship behaviors^43,44^. Using this approach, we conducted experiments to test the hypothesis that estrous state regulates female courtship USV production in inbred female mice and in outbred wild-derived female mice.

## Results

### Generation of male mice that are muted for USV production

To generate male mice that were muted for USV production, the caudal PAG of TRAP2;Ai14 male mice was injected bilaterally with a virus driving the Cre-dependent expression of caspase (AAV-FLEX-taCasp3-TEVp). Two weeks following viral injection (day 14), males were given a 30-minute courtship interaction to elicit USV production and then received IP injections of 4-hydroxytamoxifen, which enables the transient expression of Cre recombinase in recently active neurons, hence permitting the expression of caspase in PAG neurons that are required for USV production^43,44^ (Fig. 1A). Ten days later (day 24), males were tested again in courtship trials with females to evaluate USV production and non-vocal courtship behaviors. As in previous studies, this approach dramatically reduced male USV production without affecting rates of non-vocal courtship behaviors (Fig. 1B-C; p < 0.001 for change in total USVs, exact Wilcoxon rank sum test; p > 0.05 for changes in rates of non-vocal behaviors, exact Wilcoxon rank sum tests; see Table S1 for full details of statistical analyses; female estrous state was not tracked in these trials).

**Figure 1.**
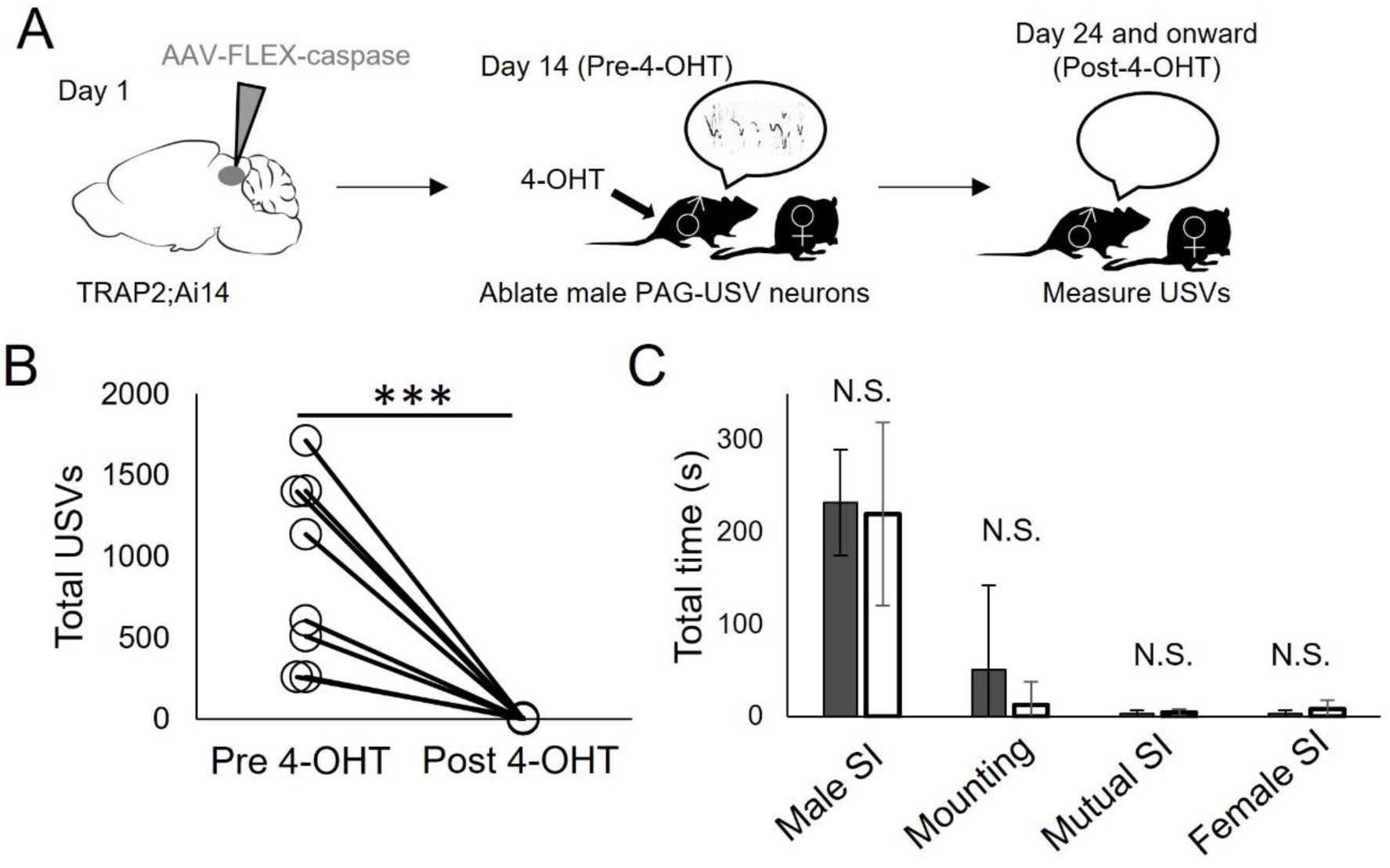
Generation of male mice that are muted for USV production. (A) Experimental timeline for TRAP2-mediated ablation of PAG neurons essential for USV production (PAG-USV neurons). (B) Total USVs recorded during courtship trials pre-vs. post-4-OHT treatment. Because pre-4-OHT trials (30 min) were longer than subsequent post-4-OHT courtship trials (10 min), total USVs produced in the first 10 minutes of pre-4-OHT trials were analyzed. (C) Same as (B), for time spent in male social investigation (SI) of the female, male mounting of the female, mutual SI, and female SI of the male. Error bars indicate standard deviation.

### Experiment 1: USV rates are low during interactions between group-housed B6 females and muted B6 males, independent of female estrous state

In Experiment 1, group-housed female mice (n = 7, 8-12 weeks-old at the start of experiment, all on a C57BL/6J background) were given a daily, 10-minute-long courtship interaction with a muted male, and female estrous state was tracked via vaginal cytology after each courtship trial. In this design, females interacted with different muted males on different days (Fig. 2A, top; n = 6 muted males were used Experiment 1). We opted for relatively short social interactions for two reasons. First, because we planned to measure behavior longitudinally from subject females, we wanted to avoid intromission and/or successful copulation during courtship trials. Second, microphone array studies have reported that female mice produce USVs during appetitive courtship behaviors, including face-to-face sniffing^35^ and when being chased by males^32^, and these behaviors occur at high rates during 10-minute courtship trials. Similarly, studies in wild-derived mice have reported that females produce USVs when interacting with males across a perforated barrier^27,36^. We performed a pair-wise comparison of total USVs between the first courtship trial in which each female was sexually receptive (proestrus/estrus, see Methods) and the first courtship trial in which each female was sexually non-receptive (metestrus/diestrus; see Figs. S1-S3 for vaginal cytology for Experiments 1-3). This analysis revealed that USV rates were overall low, and there was no significant effect of female estrous state on total USVs (Fig. 2B; mean non-receptive USVs = 0.0 ± 0.0, mean receptive USVs = 2.0 ± 4.8; p > 0.05, exact Wilcoxon rank sum test). Consistent with prior work, there was no significant effect of female estrous state on time spent by males engaged in social investigation or mounting of the female, nor on time spent by females investigating males^45^ (Fig. S4; p > 0.05 for all), and no intromissions were observed.

**Figure 2.**
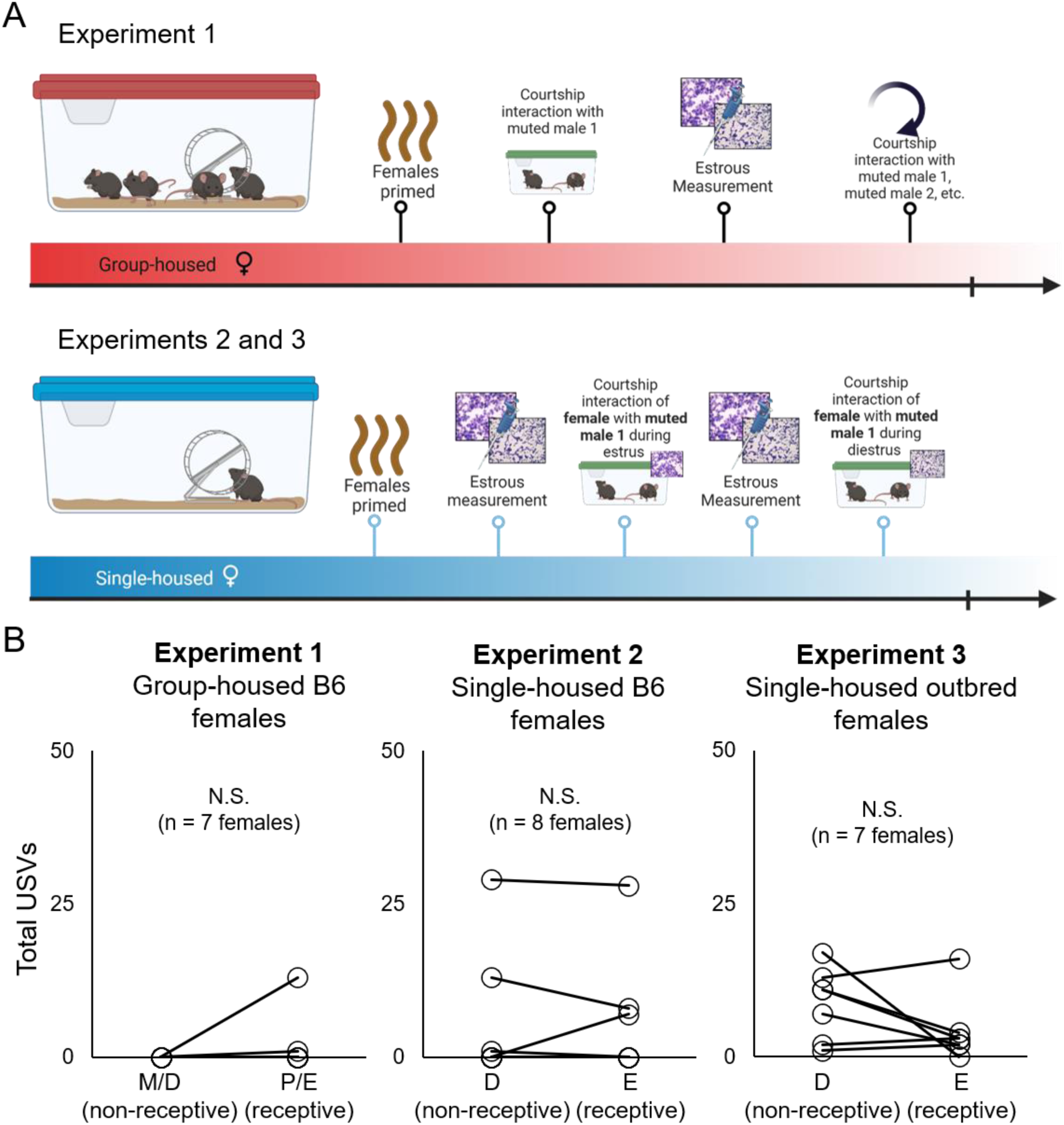
USV rates are low during interactions between female mice and muted males, independent of female estrous state. (A) Top panel shows experimental design and timeline for Experiment 1, and bottom panel shows experimental design and timeline for Experiments 2-3. (B) Total USVs are shown for courtship trials between muted males and group-housed B6 females (left), single-housed B6 females (middle), and single-housed outbred wild-derived females (right) according to female estrous state. M, metestrus; D, diestrus; P, proestrus; E; estrus.

To confirm that these USV rates are lower than what would be predicted if females produced 5-18% of total USVs in courtship trials, we calculated 18% and 5% of total USVs detected in the first 10 minutes of each pre-4-OHT courtship trial between TRAP2;Ai14 males and females. We note that the observed total USVs are lower than either of these predicted ranges (Fig. 3; 18% of total pre-4-OHT USVs = 163.7 ± 102.7; 5% of total pre-4-OHT USVs = 48.5 ± 28.5). In summary, group-housed females produce negligible rates of USVs during interactions with muted males, and these rates are not modulated by female estrous state.

**Figure 3.**
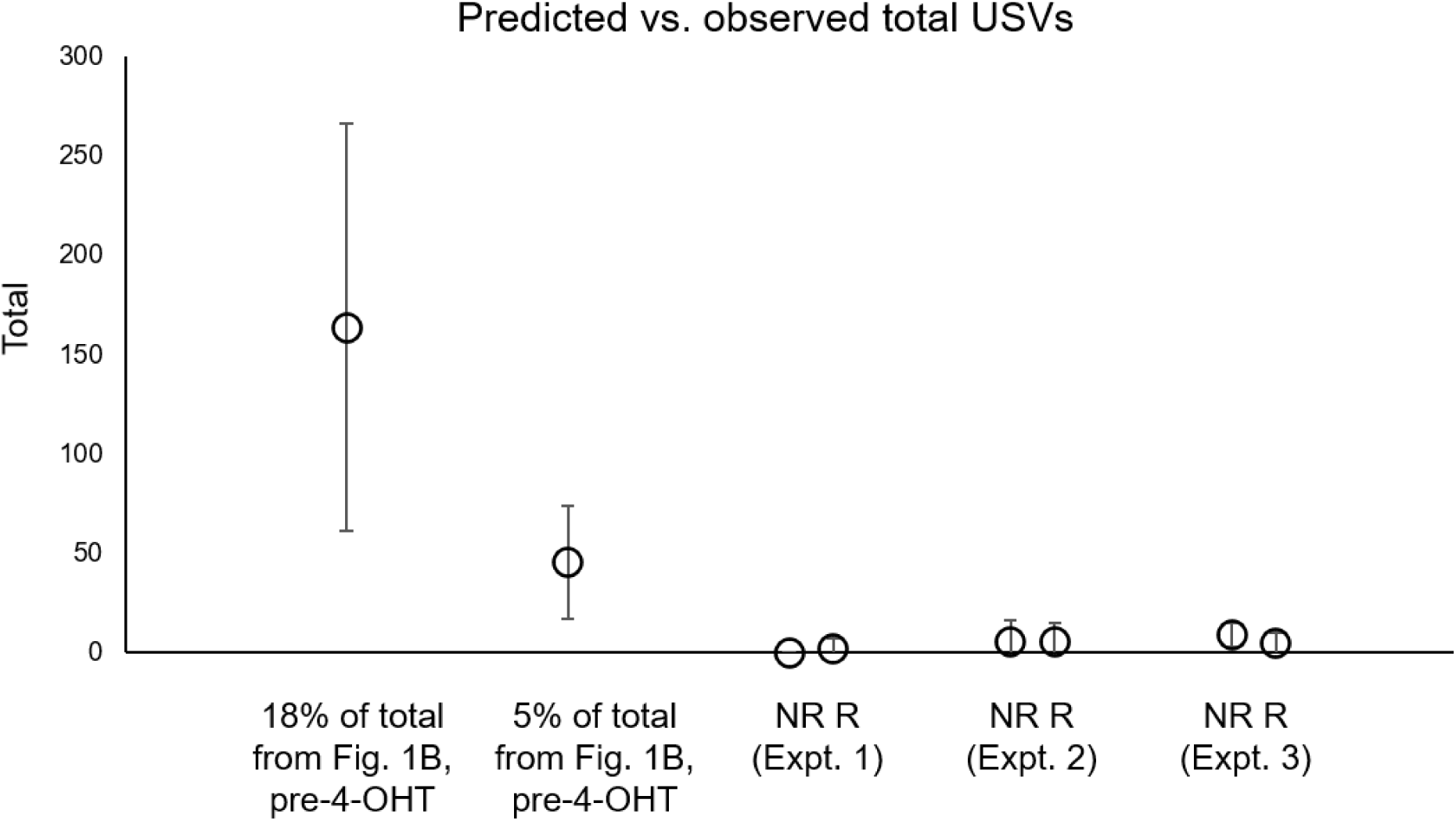
Predicted vs. observed total USVs. Predicted USVs in courtship trials with females and muted males were generated by calculating 18% and 5% of total USVs recorded in the first 10 minutes of pre-4-OHT courtship trials between TRAP2;Ai14 males and females. These predicted rates are plotted alongside the observed total USVs in Experiments 1-3. NR, non-receptive courtship trials; R, receptive courtship trials. Error bars indicate standard deviation.

### Experiment 2: USV rates are low during interactions between single-housed B6 females and muted B6 males, independent of female estrous state

In some microphone array studies that reported female USV production during mixed-sex interactions, female mice were single-housed for at least 2 weeks prior to behavioral measurements^32,34,41,42^. To assess the potential effect of housing status on female courtship USV production, we recorded USV rates in Experiment 2 during courtship trials between single-housed B6 female subjects (n = 8) and males that had been muted for USV production (n = 2 males, distinct from those used in Experiment 1). Females were single-housed for 14-18 days prior to the start of behavioral measurements, and estrous state was assessed daily starting 3 days prior to the start of behavioral measurements. In Experiment 2, we employed a fully repeated measures design in which each female was given a 10-minute courtship trial with a muted male on the first day that she was sexually receptive (defined in Experiment 2 as the first day in estrus) and then was given a second courtship trial with the same muted male on the next subsequent day that she was non-receptive (Fig. 2A, bottom; defined as the first day in diestrus). As in Experiment 1, USV rates were overall low, and there was no significant effect of female estrous state on total USVs (Fig. 2B; mean total estrus USVs = 5.4 ± 10.6, mean total diestrus USVs = 5.4 ± 9.8; p > 0.05, exact Wilcoxon rank sum test). Also as in Experiment 1, we did not find any significant effect of female estrous state on time spent in male social investigation of the female, male mounting, or female social investigation of the male (Fig. S4; p > 0.05 for all).

Because the courtship trials in both Experiments 1 and 2 took place in male home cages, we also tested a smaller cohort of single-housed B6 females in courtship trials that occurred within the female’s home cage. Total USVs in these trials were also low, independent of female estrous state (n = 4 females; mean total estrus USVs = 1.3 ± 2.5; mean total diestrus USVs = 3.8 ± 3.8). These findings indicate that single-housed females produce negligible rates of USVs during interactions with muted males and that these USV rates are not modulated by female estrous state.

### Experiment 3: USV rates are low during interactions between single-housed outbred females and muted B6 males, independent of female estrous state

Recent work comparing mixed-sex populations of C57BL/6J mice to mixed-sex populations of wild-derived mice in seminatural enclosures found that inbred and wild-derived females differ dramatically in their social behaviors^46^. Moreover, studies using a split-cage paradigm to record USVs from mice interacting across a perforated barrier report that wild-derived female mice produce USVs during such interactions with males^27,36^. Given these findings, one idea is that female courtship USV production is low in inbred female but may be more robust in wild-derived females. In Experiment 3, we examined whether our findings in inbred females could be extended to wild-derived females by recording total USVs during 10-minute courtship trials between outbred wild-derived single-housed female subjects^46^ (n = 7) and males that had been muted for USV production (n = 2; same muted males and experimental design as Experiment 2). As in Experiments 1 and 2, we found that USV rates were overall low and that there was no significant effect of female estrous state on total USVs (Fig. 2C; mean non-receptive USVs = 8.9 ± 5.8, mean receptive USVs = 4.3 ± 5.3; p > 0.05, exact Wilcoxon rank sum test). There was also no effect of female estrous state on time spent in male social investigation of the female, male mounting, or female social investigation of the male (Fig. S4; p > 0.05 for all). Because B6 females are known to produce high rates of USVs during same-sex interactions^20,22,23^, particularly following single-housing^24,25^, we also confirmed that single-housed outbred wild-derived females produced high rates of USVs when paired together in round-robin same-sex interactions (mean total USVs = 900.4 ± 493.4, from n = 9, 10-minute-long female-female trials). These experiments show that although outbred wild-derived females produce high rates of USVs during same-sex interactions, they produce low USV rates during courtship interactions with muted males, regardless of estrous state.

### Female USV production vs. residual USV production by muted males

Although our experiments are designed to facilitate the detection of female USVs, it is possible that some or all of the USVs in our trials are attributable to a small amount of residual USV production by the muted males. To evaluate this idea, we compared the temporal overlap between USVs and squeaks during courtship trials. Female mice are well known to produce squeaks during courtship interactions with males, both when sexually receptive and non-receptive^47–49^. Because USVs and squeaks are produced via different phonation mechanisms, a mouse cannot produce both vocalization types simultaneously. Therefore, any USV that overlaps temporally with a squeak can be attributed to the male. In Experiment 1, we found that 1 of 14 USVs (7.1%) overlapped with a squeak. In Experiments 2 and 3, the numbers of overlapping USVs were 6 of 86 (7.0%) and 5 of 92 (5.4%), respectively. If we broaden our consideration to USVs that occur within 0.5s of a squeak, the counts of USVs that occur overlapping with or adjacent to a squeak are 2 of 14 (14.3%, Experiment 1), 9 of 86 (10.5%, Experiment 2), and 11/92 (12.0%, Experiment 3). These analyses support the idea that at least some of the USVs recorded in our courtship trials represent low rates of residual USVs produced by muted males.

## Discussion

In the current study, we tested the hypothesis that sexual receptivity promotes female USV production during courtship interactions with males. To facilitate the detection of female USVs, females were paired with males that were muted for USV production via ablation of midbrain neurons whose activity is essential for USV production. Contrary to our hypothesis, we detected low (near-zero) rates of USVs during courtship interactions between females and muted males, and these rates did not vary according to female estrous state. This finding held true for group-housed B6 background female mice (Experiment 1), single-housed B6 background female mice (Experiment 2), and single-housed outbred wild-derived female mice (Experiment 3).

How can we reconcile these near-zero USV rates with reports that females produce a portion (5-18%) of total USVs during courtship interactions? One possibility is that female mice only produce appreciable rates of USVs during interactions with vocal male partners. There are three interesting considerations related to this idea. First, in females of other rodent species that are known to produce courtship USVs, female courtship USV production is regulated by estrous state but is not contingent on male USV production. For example, female hamsters produce USVs both during as well as following interactions with males^39^ and during investigation of male bedding and anesthetized males^37^. Female rats produce USVs during interactions with male partners, even those that have been devocalized via transection of the recurrent laryngeal nerve^50^, and female USV rates during interactions with devocalized males are higher during proestrus/estrus than during diestrus^40^. Therefore, if courtship USV production in female mice is contingent on male vocal production, this behavioral feature would represent a notable divergence from other rodent species. Second, female mice produce USVs when interacting with other females^20,22–25^, and they continue to produce USVs even when interacting with females that have been muted for USV production via the same strategy employed in the current study^44^. Thus, the requirement for a vocal partner would also need to be context-dependent (i.e., only exhibited by females during courtship interactions but not during same-sex interactions). Third, male mice produce courtship USVs when interacting with devocalized female partners^14,17^, and hearing vs. deafened male mice produce courtship USVs at identical rates and with similar acoustic features during interactions with females^51^. Given that male mouse courtship USV production does not appear to require auditory information of any type, female courtship USV production that is contingent on partner vocal production would also represent a sex-specific trait in mice.

A second (and in our opinion, more likely) possibility is that female mice vocalize at negligible rates during courtship interactions, even those with intact males, and that some microphone array studies have overestimated female courtship USV production. Courtship USVs are produced when males and females are in close proximity and often when they are moving, which presents challenges to accurate localization and assignment of USVs to individual signalers. Recent studies have improved USV localization accuracy from ∼38 mm^32^ to ∼14 mm^30^ to ∼4 mm^35^, which has enabled high-confidence assignment of a larger proportion of total USVs to individual signalers and would in turn be expected to reduce errors in USV assignment. It is notable that alongside these improvements in localization accuracy, the percentage of USVs attributed to female signalers during courtship interactions has generally decreased (from 18%^32,34^ to 16%^30^ to 10%^33^ to 5%^35^). In the most recent of such studies, Sterling et al., 2023 note further that when consideration is limited to USVs produced when male and female snouts are separated by a distance of > 50 mm, the percentage of USVs assigned to the female drops from ∼5% to ∼1%. Aside from differences in estimates of female USV production resulting from differences in localization accuracy, however, it is important to note that the use of non-receptive females (likely the case in studies that did not track estrous state^30,33,35^) and the use of smaller, less natural behavioral set-ups (such as elevated platforms^35^) may tend to suppress any female USV production and could also contribute to differences in results between these studies.

Distinguishing between these two possibilities will require methodologies that can unambiguously assign the vast majority of USVs to a signaler, regardless of inter-mouse distances and movement speeds. One possibility is that measurements of respiration from freely moving male mice could be applied to assign USVs that align with the male’s exhalations as originating from the male, while any USVs misaligned to male respiration would be assigned to the female. Such a method was applied in a recent study to examine the relationship of USV production to sniffing in rats and mice^52^. Notably, in the subset of their experiments conducted in male mice interacting with females, the authors report that all recorded USVs were aligned to the male respiratory cycle^52^. Alternatively, electrophysiological recordings from the PAG or from vocalization-related hindbrain neurons that regulate USV production could in principle be used to assign vocalizations to individual signalers with high fidelity during free interactions between females and males. If such approaches provide compelling evidence for female courtship USV production, subsequent studies could revisit the question of whether this behavior is regulated by female estrous state.

Given that previous studies in wild-derived mice reported that females produce USVs when interacting with males across a perforated barrier^27,36^, we were surprised to find such low rates of USV production in courtship trials between muted males and outbred wild-derived females. Although the wild-derived females used in our study mate readily with B6 males, we cannot exclude the possibility that their courtship behaviors, including potential USV production, may differ in interactions with wild-derived and B6 males. Whether female vocal behavior differs between inbred and wild-derived mice, during courtship interactions as well as during same-sex interactions^26,27,36^, remains an important and ongoing topic of study.

## Materials and Methods

### Ethics Statement

All experiments and procedures were conducted according to protocols approved by the Cornell University Institutional Animal Care and Use Committee (protocol #2020-001).

### Subject Details

Male and female mice were housed with their siblings and both parents until weaning at postnatal day 21. All males used in the study were > 8 weeks of age, and females used in the study were 8-12 weeks of age at the time of behavioral measurements. All male mice used in the study were TRAP;Ai14, which were generated by crossing TRAP2 (Jackson Laboratories, 030323) with Ai14 (Jackson Laboratories, 007914). Female laboratory mice used in the study were from various strains, all on a C57BL/6J background (TRAP2 - 030323; Ai14 - 007914, TRAP2;Ai14, or C57BL/6J - 000664). Outbred wild-derived female mice were obtained from the laboratory of Dr. Michael Sheehan, and the progenitors of these mice originated from Saratoga Springs, NY^46^. Outbred females were allowed to acclimate for 2 weeks after being transferred between colony rooms prior to the start of behavioral measurements. Mice were kept on a 12h:12h reversed light/dark cycle and given ad libitum food and water for the duration of the experiment.

### Drug preparation

4-hydroxytamoxifen (4-OHT, HelloBio, HB6040) was dissolved at 20 mg/mL in ethanol by shaking at 37°C and was then aliquoted (75 uL) and stored at −20°C. Before use, 4-OHT was redissolved in ethanol by shaking at 37°C and filtered corn oil was added (Sigma, C8267, 150 uL). Ethanol was then evaporated by vacuum under centrifugation to give a final concentration of 10 mg/mL, and 4-OHT solution was used on the same day it was prepared.

### Ablation of PAG-USV neurons in male mice

To express caspase in PAG-USV neurons, male mice first received bilateral injections of virus (AAV2/5-ef1α-FLEX-taCasp3-TEVp, Addgene, 45580) into the caudolateral PAG (day 0). The final injection coordinates for caudolateral PAG were: AP = −4.7 mm, ML = 0.5 mm, DV = 1.75 mm. Virus was pressure-injected with a Nanoject III (Drummond) at a rate of 5 nL every 20 seconds. A total volume of 250 nL of virus was injected into each side of the PAG. Eleven days later, males were single-housed for 3 days. On day 14, males were given a 30-minute courtship interaction with a novel, group-housed female visitor to elicit USV production. Following the social interaction, males received I.P. injections of 4-OHT (50 mg/kg) to enable expression of caspase in PAG-USV neurons. Males remained single-housed throughout the duration of the experiment to promote high levels of social motivation during subsequent courtship trials with subject females^24,25^. Males were used in subsequent courtship trials following a wait time of at least 10 days post-4-OHT treatment.

### Post-hoc visualization of viral labeling

Mice were deeply anesthetized with isoflurane and transcardially perfused with ice-cold 4% paraformaldehyde in 0.1 M phosphate buffer, pH 7.4 (4% PFA). Dissected brains were post-fixed overnight in 4% PFA at 4°C, cryoprotected in a 30% sucrose solution in PBS at 4°C for 48 hours, frozen in embedding medium (Surgipath, VWR), and stored at −80°C until sectioning. Brains were cut into 80 µm coronal sections on a cryostat, rinsed 3 x 10 minutes in PBS, and processed at 4°C with NeuroTrace (1:500, Invitrogen) in PBS containing 0.3% Triton-X. Tissue sections were rinsed again 3 x 10 minutes in PBS, mounted on slides, and coverslipped with Fluoromount-G (Southern Biotech). After drying, slides were imaged with a 10x objective on a Zeiss LSM900 confocal laser scanning microscope. Because expression of the caspase virus cannot be directly visualized by looking for expression of a fluorescent tag, we took the following approach to assess the spread of the caspase virus. In TRAP2;Ai14 males that are treated with 4-OHT following a social encounter, neurons throughout the brain that upregulate Fos during the social encounter will be labeled with tdTomato. Because all TRAPed, caspase-expressing PAG-USV neurons are subsequently ablated, viral spread was examined by assessing the absence of tdTomato labeling in the PAG of TRAP2;Ai14 males^44^. N = 2 males with viral spread that did not fully and bilaterally cover the caudolateral PAG were excluded from the study.

### Measurements of female estrous state

Vaginal cytology samples were collected by vaginal lavage with PBS (50 μL). Samples were pipetted directly onto slides and allowed to dry. After drying, slides were stained with Jorvet Dip Quick Stain (Jorgensen Laboratories) according to the manufacturer instructions. Stained slides were imaged at 20x magnification on a light microscope, and estrous state was determined by cell proportions as outlined previously^53^. In Experiment 1, females in proestrus and estrus were classified as receptive, and females in metestrus and diestrus were classified as non-receptive. In Experiments 2 and 3, behavioral tests were performed only on days during which females were in estrus or in diestrus. Cytology images for Experiments 1-3 are included in Figs. S1-S3.

### Experiment 1 design

Males (n = 6) used in Experiment 1 were muted for USV production via ablation of PAG-USV neurons as described above and in previous work^43,44^. Female subjects used in Experiment 1 (n = 7) were group-housed with same-sex siblings. Three days prior to the start of courtship trials, male bedding was added to the female home cage. Three days later, the subject female was placed in the home cage of a muted male inside a sound-attenuating chamber equipped with an infrared (IR) light source, a webcam (IR filter removed to enable video recording under IR lighting conditions), and an ultrasonic microphone. A custom lid was placed on the cage to allow video and audio recordings, and vocal and non-vocal behaviors were recorded for 10 minutes. Female estrous state was measured as described above following the courtship trial, and the female was then returned to her home cage with her same-sex siblings. These courtship trials and estrous state measurements were repeated daily for each subject female until we obtained one courtship trial in which the female was sexually receptive (proestrus/estrus) and another courtship trial in which the female was sexually non-receptive (metestrus/diestrus). The first courtship trial of each type obtained for a given female was used for subsequent analyses. In Experiment 1, females were paired with different muted males in different courtship trials.

### Experiment 2 design

Experiment 2 was modified from the design of Experiment 1 in the following ways. First, females used in Experiment 2 (n = 8) were single-housed for at least 2 weeks (14-18 days) prior to the start of courtship trials. Second, the priming procedure was modified to include a 10-minute social interaction with a male in addition to the placement of male bedding in the female’s home cage. Priming occurred 3 days prior to the start of courtship trials. Third, female estrous state was measured daily starting 3 days post-priming, and the first courtship trial was conducted on the first day that the female was in estrus (trial began ∼1.5 hours after the estrous measurement). The second courtship trial was then conducted on the first subsequent day that the female was in diestrus. Finally, Experiment 2 employed a fully repeated measures design in which a given female was tested with the same muted male twice, once while sexually receptive and once while non-receptive. Measurements of vocal and non-vocal social behaviors during courtship trials were collected as in Experiment 1. Males (n = 2) used in Experiment 2 were muted for USV production via ablation of PAG-USV neurons but were distinct from the males used in Experiment 1.

### Experiment 3 design

Experiment 3 was conducted similarly to Experiment 2, except that the females (n = 7) used were outbred wild-derived females obtained from the laboratory of Dr. Michael Sheehan^46^. Outbred females were allowed to acclimate for 2 weeks after being transferred between colony rooms prior to the start of behavioral measurements. Following the 2-week acclimation period, all experimental procedures and measurements were conducted as in Experiment 2. The same muted males (n = 2) were used in courtship trials in Experiments 2 and 3.

### USV recording and analyses

USVs were recorded with an ultrasonic microphone (Avisoft, CMPA/CM16), amplified (Presonus TubePreV2), and digitized at 250 kHz (Spike 7, CED). Because USV rates were low across all three experiments and because low amplitude USVs are often missed by automated USV detection codes, USVs were manually scored by visual observation of spectrograms from courtship trials by a trained observed (C.A.M.).

### Analyses of non-vocal social behaviors

BORIS software was used by trained observers to score and record non-vocal social behaviors from video recordings of pairs of interacting males and females. The following behaviors were recorded: male social investigation of the female (sniffing, following, or chasing), female social investigation of the male, mutual social investigation, and male mounting of the female. Intromission, fighting, and instances of female subjects mounting males were not observed in any trial.

### Statistics

The Shapiro-Wilk test was used to evaluate the normality of data distributions. Parametric, two-sided statistical comparisons were used for normally distributed data, and non-parametric two-sided comparisons were used for non-normally distributed data (alpha = 0.05). No statistical methods were used to pre-determine sample size. Full details of statistical analyses are included in Table S1.

## Data Availability

The datasets generated during the current study will be made available publicly and without restriction from Cornell eCommons, and this section will be updated with a persistent DOI upon acceptance of the manuscript for publication.

## Acknowledgements

Thanks to Mike Sheehan for providing us with outbred wild-derived females and for helpful discussions of the project. Thanks to Caitlin Miller for assistance with estrous staining. Thanks also to Frank Drake for his excellent mouse husbandry.

## Author contributions

C.A.M. and P.Z. performed viral injections on male mice. C.A.M. performed behavioral experiments and vaginal cytology. C.A.M., P.Z., J.K., and K.A.T. analyzed data. C.A.M. and K.A.T. wrote the manuscript, and all authors approved the final manuscript.

## Complete interests

The authors declare no competing interests.

