## Supplemental Information for "Rates of female mouse ultrasonic vocalizations are low and are not modulated by estrous state during interactions with muted males"

**Supplemental Information for Malone et al.**

Figures S1-S4

Table S1

Figure S1: Vaginal cytology for Experiment 1

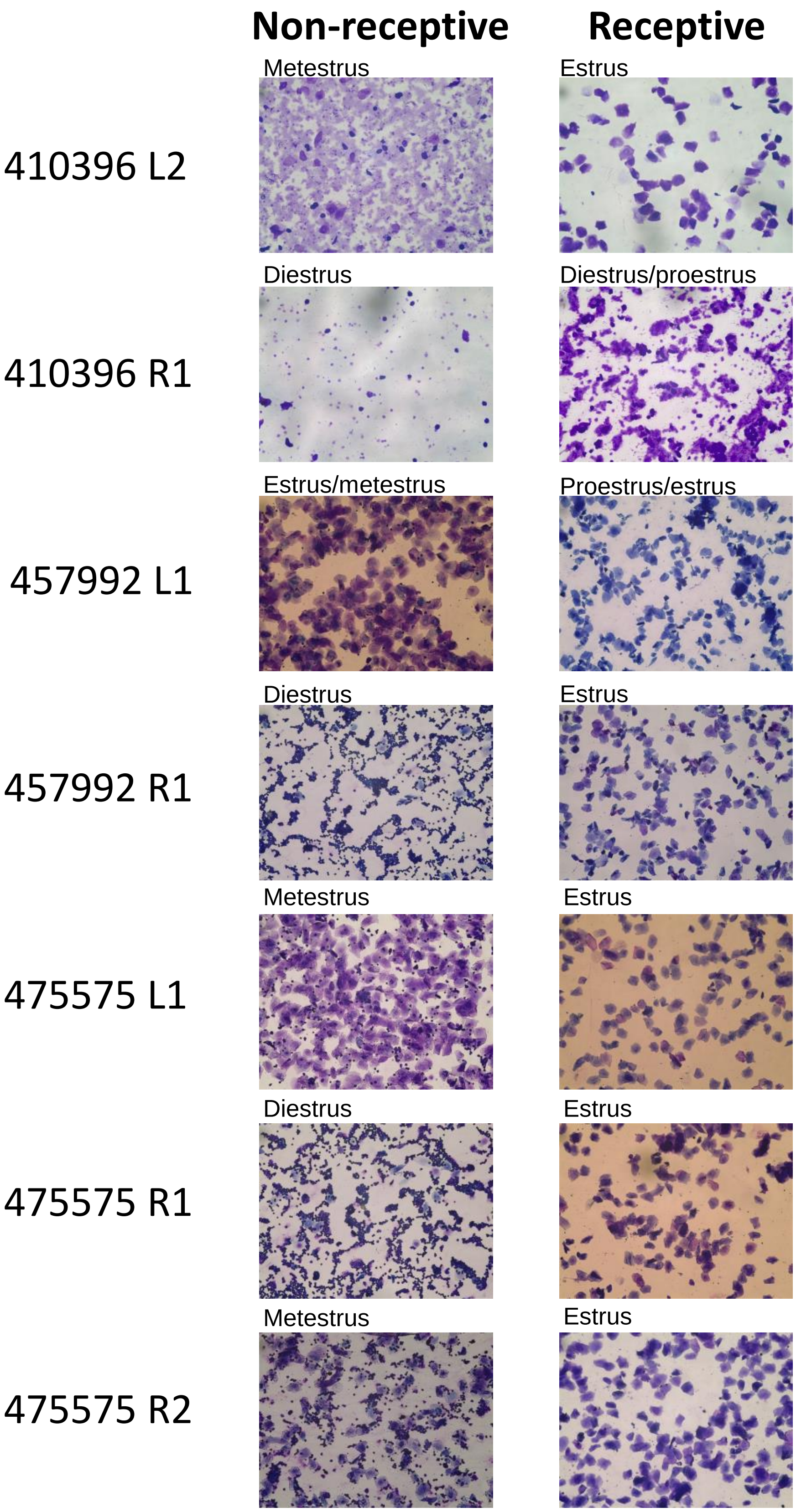

### Figure S2: Vaginal cytology for Experiment 2

Disestrus | Estrus

504705 NT
504705 R1
511167 L1
511167 NT
516389 L1
516389 NT
520391 L1
520391 R1

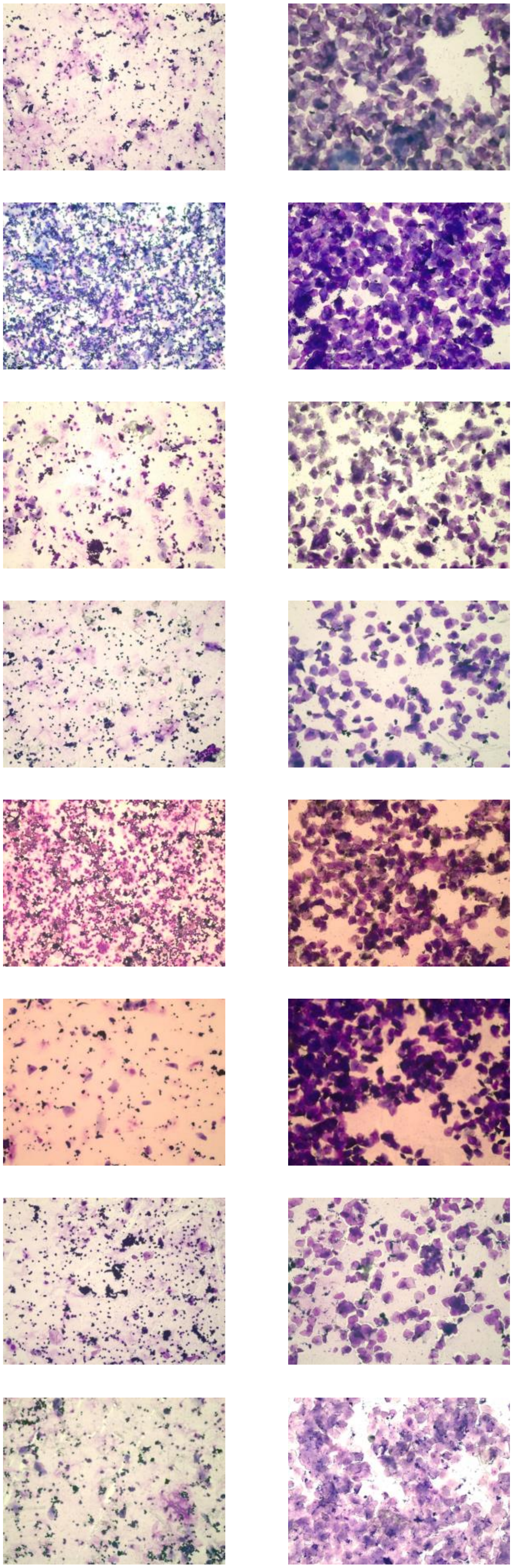

Figure S3: Vaginal cytology for Experiment 3

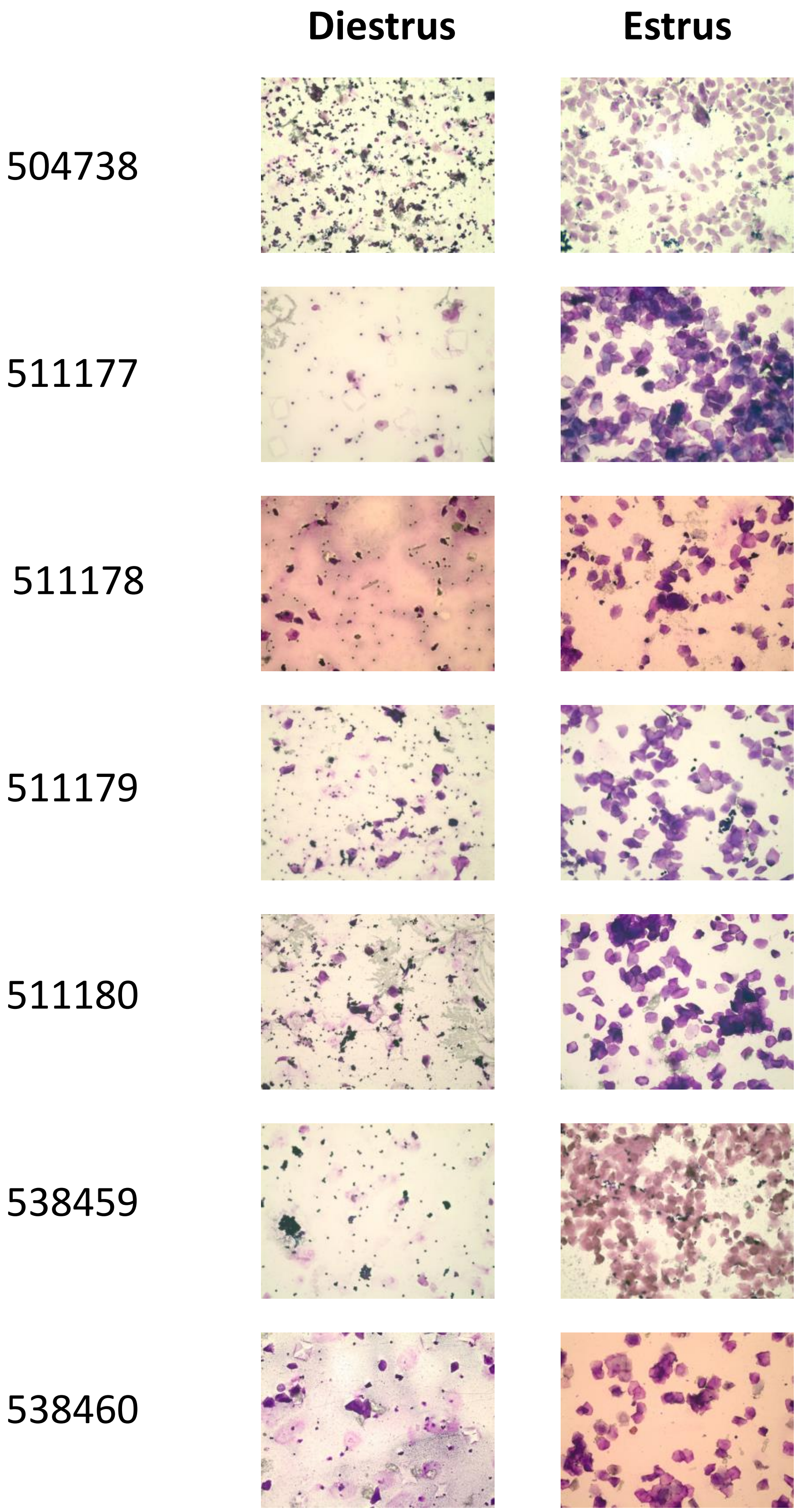

### Figure S4: Quantification of non-USV social behaviors

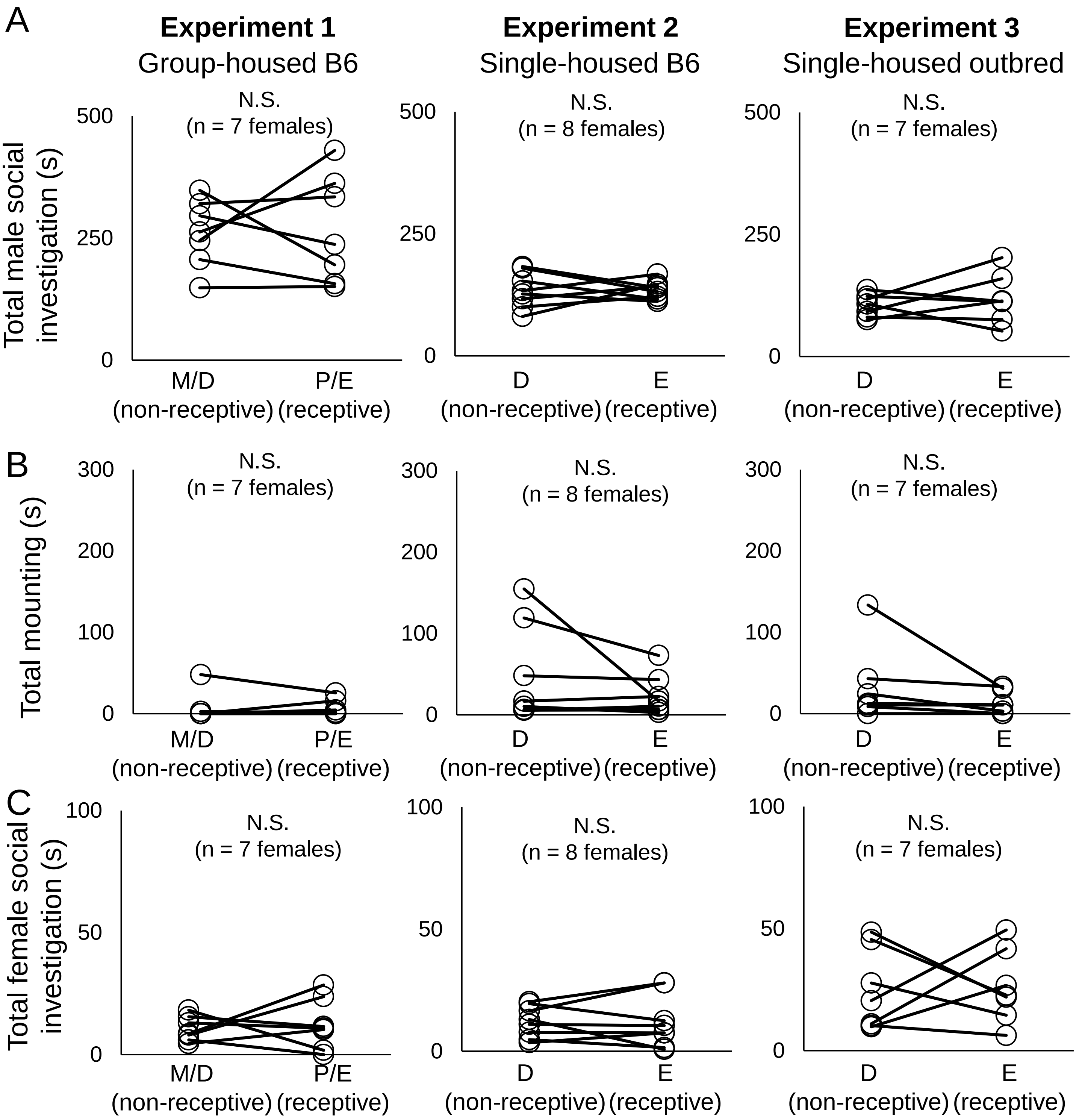

**Table S1.** Details of statistical analyses

| **Figure, comparison, and statistical test** | **Group means +/- SD** | **Test results** |
|---|---|---|
| Fig. 1B: total USVs, pre- vs. post-4-OHT<br>• Exact Wilcoxon rank sum test | Pre-4-OHT = 909.6 ± 570.6, N = 8<br><br>Post-4-OHT = 1.0 ± 2.8, N = 8 | W = 81, **P < 0.01** |
| Fig. 1C: male social investigation time, pre- vs. post-4-OHT<br>• Exact Wilcoxon rank sum test | Pre-4-OHT = 232.0 ± 55.8, N = 8<br><br>Post-4-OHT = 219.8 ± 99.4, N = 8 | W = 39, P = 0.51 |
| Fig. 1C: male mounting time, pre- vs. post-4-OHT<br>• Exact Wilcoxon rank sum test | Pre-4-OHT = 50.8 ± 91.4, N = 8<br><br>Post-4-OHT = 12.4 ± 25.0, N = 8 | W = 39.5, P = 0.44 |
| Fig. 1C: mutual social investigation time, pre- vs. post-4-OHT<br>• Exact Wilcoxon rank sum test | Pre-4-OHT = 3.1 ± 3.7, N = 8<br><br>Post-4-OHT = 4.4 ± 3.5, N = 8 | W = 25, P = 0.50 |
| Fig. 1C: female social investigation time, pre- vs. post-4-OHT<br>• Exact Wilcoxon rank sum test | Pre-4-OHT = 2.9 ± 3.3, N = 8<br><br>Post-4-OHT = 8.0 ± 9.9, N = 8 | W = 24, P = 0.43 |
| Fig. 2B, left: total USVs, Experiment 1<br>• Exact Wilcoxon rank sum test | Non-receptive = 0.0 ± 0.0, N = 7<br><br>Receptive = 2.0 ± 4.9, N = 7 | W = 17.5, P = 0.46 |
| Fig. 2B, middle: total USVs, Experiment 2<br>• Exact Wilcoxon rank sum test | Non-receptive = 5.4 ± 10.6, N = 8<br><br>Receptive = 5.4 ± 9.8, N = 8 | W = 32.5, P = 1 |
| Fig. 2B, right: total USVs, Experiment 3<br>• Exact Wilcoxon rank sum test | Non-receptive = 8.9 ± 5.9, N = 7<br><br>Receptive = 4.3 ± 5.3, N = 7 | W = 34, P = 0.24 |

| | | |
|---|---|---|
| Fig. S4A, left: total male social investigation, Experiment 1<br>• Paired t-test | Non-receptive =<br>261.0 ± 68.7, N = 7<br><br>Receptive =<br>266.7 ± 109.4, N = 7 | T(6) = -0.14, P = 0.89 |
| Fig. S4A, middle: total male social investigation, Experiment 2<br>• Paired t-test | Non-receptive =<br>134.2 ± 36.3, N = 8<br><br>Receptive =<br>134.6 ± 18.4, N = 8 | T(7) = -0.03, P = 0.98 |
| Fig. S4A, right: total male social investigation, Experiment 3<br>• Paired t-test | Non-receptive =<br>104.7 ± 22.9, N = 7<br><br>Receptive =<br>118.4 ± 50.1, N = 7 | T(6) = -0.71, P = 0.50 |
| Fig. S4B, left: total male mounting, Experiment 1<br>• Exact Wilcoxon rank sum test | Non-receptive =<br>7.2 ± 18.0, N = 7<br><br>Receptive =<br>6.8 ± 10.0, N = 7 | W = 18.5, P = 0.46 |
| Fig. S4B, middle: total male mounting, Experiment 2<br>• Exact Wilcoxon rank sum test | Non-receptive =<br>45.8 ± 58.7, N = 8<br><br>Receptive =<br>22.7 ± 24.1, N = 8 | W = 34, P = 0.88 |
| Fig. S4B, right: total male mounting, Experiment 3<br>• Exact Wilcoxon rank sum test | Non-receptive =<br>33.2 ± 46.3, N = 7<br><br>Receptive =<br>12.7 ± 14.0, N = 7 | W = 32, P = 0.38 |
| Fig. S4C, left: total female social investigation, Experiment 1<br>• Paired t-test | Non-receptive =<br>10.5 ± 5.1, N = 7<br><br>Receptive =<br>12.4 ± 10.5, N = 7 | T(6) = -0.38, P = 0.72 |
| Fig. S4C, middle: total female social investigation, Experiment 2<br>• Paired t-test | Non-receptive =<br>12.0 ± 6.4, N = 8<br><br>Receptive =<br>12.0 ± 10.6, N = 8 | T(7) = 0, P = 1.0 |

| | | |
|---|---|---|
| Fig. S4C, right: total female social investigation, Experiment 3<br>• Paired t-test | Non-receptive = 24.8 ± 16.6, N = 7<br><br>Receptive = 26.2 ± 15.0, N = 7 | T(6) = -0.17, P = 0.87 |
